# Machine Learning Approaches for Skin Neoplasm Diagnosis

**DOI:** 10.1101/2024.05.12.593773

**Authors:** Abu Asaduzzaman, Christian C. Thompson, Md J. Uddin

## Abstract

Approaches for skin neoplasm diagnosis include physical exam, skin biopsy, lab tests of biopsy samples, and image analyses. These approaches often involve error-prone and time-consuming processes. Recent studies show that machine learning has promises to effectively classify skin images into different classes such as melanoma and melanocytic nevi. In this work, we investigate machine learning approaches to enhance the performance of computer-aided diagnosis (CADx) systems to diagnose skin diseases. In the proposed CADx system, generative adversarial network (GAN) is used to identify (and remove) fake images. Exploratory data analysis (EDA) is applied to normalize the original dataset for preventing model overfitting. Synthetic minority over-sampling technique (SMOTE) is employed to rectify class imbalances in the original dataset. To accurately classify skin images, the following four machine learning models are utilized: linear discriminant analysis (LDA), support vector machine (SVM), convolutional neural network (CNN), and an ensemble CNN-SVM. Experimental results using the HAM10000 dataset demonstrate the ability of the machine learning models to improve CADx performance in treating skin neoplasm. Initially, the LDA, SVM, CNN, and ensemble CNN-SVM show 49%, 72%, 77%, and 79% accuracy, respectively. After applying GAN, EDA, and SMOTE, the LDA, SVM, CNN, and ensemble CNN-SVM show 76%, 83%, 87%, and 94% accuracy, respectively. We plan to explore other machine learning models and datasets in our next endeavor.

## 1. Introduction

Machine learning has emerged as a powerful tool, analyzing complex patterns and features in neoplasms to assist dermatologists in identifying malignant lesions.^1–3^ One of the primary applications of classification techniques is in computer-aided diagnostic (CADx) systems. ^4,5^ These systems typically involve training algorithms on large datasets of annotated medical images to learn the characteristic features associated with different types of skin lesions.^6,7^ Machine learning based CADx systems offer several advantages including early detection, improved accuracy, and personalized medicine. It should be noted that the machine learning based approaches need large and diverse training datasets, potential biases in data collection and annotation, interpretability issues with complex models, and regulatory considerations regarding clinical validation and deployment. Despite implementation challenges, machine learning holds great promise for the diagnosis of skin neoplasms.

The detection and diagnosis of skin diseases typically rely on the expertise of dermatologists through visual examination techniques such as the use of dermatoscopes and/or invasive procedures such as surgical biopsies.^8–11^ However, these traditional approaches are often time-consuming and expensive. To overcome these limitations, CADx systems have emerged as valuable tools for assisting in the detection of skin malignancies. An event when a CADx system incorrectly identifies a benign lesion as a cancerous lesion is called a false positive.^1,12–16^ On the other hand, when a CADx system incorrectly identifies a cancerous lesion as a benign lesion is called a false negative. ^17–19^ False positive can lead to unnecessary biopsies with many follow-up procedures, and false negative can lead to delays in treatment, making it difficult to treat. Recent studies suggest that CADx systems have demonstrated notable advancements in enhancing diagnostic accuracy by applying various machine learning concepts and techniques.

This work aims to investigate contemporary popular machine learning approaches to CADx systems for better identifying dermatological abnormalities in skin images and reduce the number of false negative and false positive readouts. This study involves the use of various classification algorithms, including LDA, SVM, CNN, and an ensemble CNN-SVM model. To enhance CADx performance, fake images are identified (and removed) using GAN. The original dataset is normalized using EDA to avoid model overfitting. Class imbalances in the original dataset is rectified using SMOTE.

This manuscript is organized as follows. Related work is reviewed in Section 2. Proposed CADx system with machine learning is described in Section 3. Experimental setup and experimental results are presented in Sections 4 and 5, respectively. Finally, this paper is concluded in Section 6.

## 2. Related Work

### 2.1 Conventional CADx Systems

Conventional CADx systems utilize computer algorithms to aid healthcare professionals in diagnosing diseases.^4,20,21^ These systems operate by furnishing clinicians with information regarding potential disease presence and by scrutinizing data derived from medical imaging modalities such as X-rays, magnetic resonance imaging (MRI), and computed tomography (CT) scans. Recognized as valuable tools, conventional CADx systems enhance the accuracy and efficiency of medical diagnoses. ^22,23^ They follow a series of steps to analyze medical images, encompassing preprocessing, segmentation, feature extraction, feature selection, and classification as illustrated in Figure 1.

**Figure 1:**
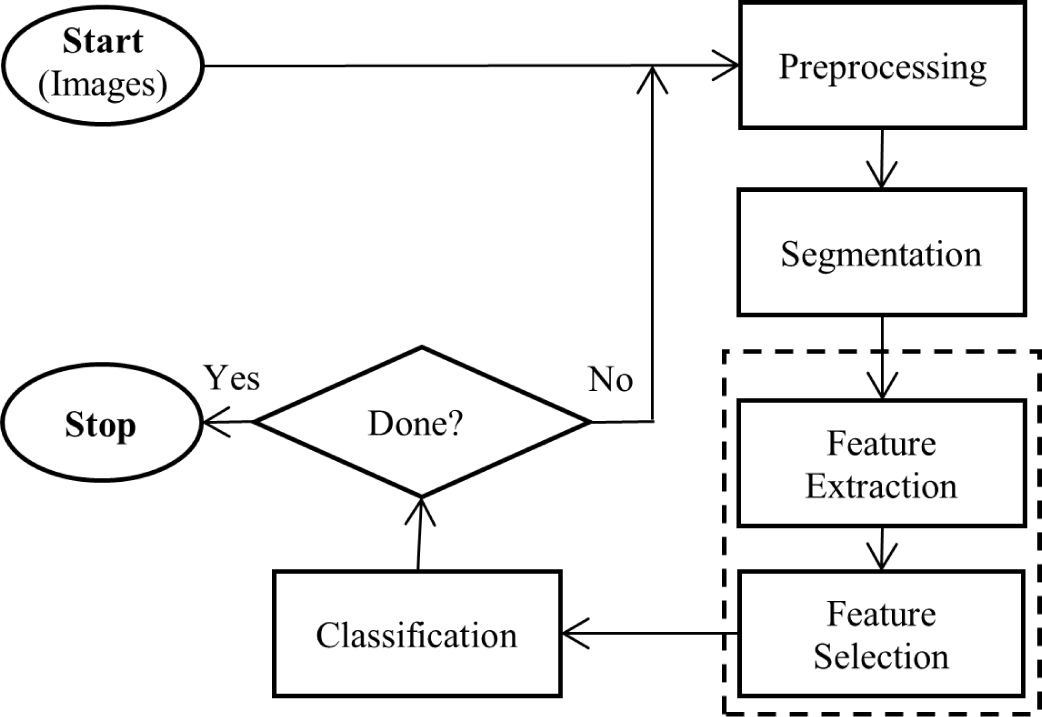
Major Steps of a conventional CADx system

The preprocessing step aims to improve the quality of the medical image by applying various techniques such as noise removal, geometric transformation, cropping, resizing, and adjusting the color balance of the images.^24,25^ The preprocessed image proceeds to the segmentation step. The segmentation step aims to identify and isolate areas of interest within the image.^26–28^ Figure 2 shows an example of the implementation of noise removal from the preprocessing stage and thresholding used in segmentation.

**Figure 2:**
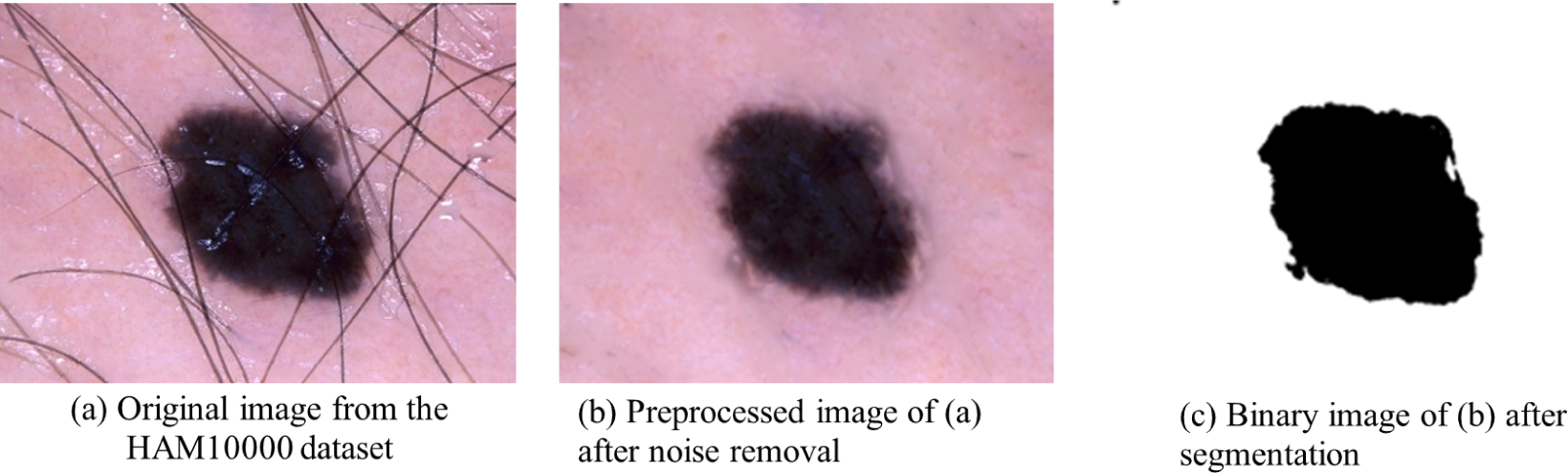
Noise Removal and Segmentation

Feature extraction and selection, two important steps, are usually grouped in CADx systems. Feature extraction involves the technique of detecting and extracting the most critical and relevant characteristics from segmented regions. ^29–31^ The goal of feature extraction is to minimize the dimensionality of the data so that it can be processed more easily and quickly while maintaining as much information as possible. Feature selection involves the process of selecting the most relevant and informative characteristics from a vast set of features derived from medical images. ^32–34^ The purpose is to minimize the dimensionality of the feature space and remove redundant characteristics that might affect CADx system performance. The important features (also known as regions of interest) are used as input in the classification step. The choice of a classification method, from three major methods as shown in Table 1, for a CADx system depends on the trade-offs between accuracy, complexity, and interpretability, as well as the specific requirements of the application at hand. Rule-based algorithms rely on explicit rules and conditions defined by experts. Statistical methods extract quantitative features from images. Machine learning models learn patterns from labeled data to make predictions.

**Table 1:** Major Classification Methods.

| **Classification Method** | **Key Applications** |
| --- | --- |
| Rule-based | Typically used for simple applications where accuracy is not critical. |
| Statistical | Typically used for applications where accuracy is important, but the system needs to be easy to understand and comprehend. |
| Machine Learning | Typically used for applications where accuracy is critical, and the system can be complex. |

In our work, we aim to improve the performance of CADx system by removing fake images, resampling data, and using an ensemble of CNN and SVM model.

### 2.2 Dataset for Studying Skin Neoplasm

In this work, we use the Human Against Machine 10000 (HAM10000) dataset, a popular one for skin neoplasm study. HAM10000 is created by Philipp Tschandl, Cliff Rosendahl, and Harald Kittler at the Medical University of Graz in Austria and is available via Kaggle website.^35–38^ HAM10000 dataset has more than 10,000 training images for detection of pigmented skin lesions with the following seven classes. Melanoma (MEL) is considered the most serious and potentially life-threatening type of skin cancer, which develops from pigment-producing cells. Basal Cell Carcinoma (BCC) is the most common type of skin cancer that typically appears as a waxy bump or lesion on the skin. Vascular Lesion (VAS) is a variety of skin conditions caused by abnormal blood vessels, including birthmarks and hemangiomas. Actinic Keratoses (AKIEC) is a pre-cancerous lesion that appears as a scaly or crusty growth and is typically caused by sun damage. Benign Keratosis-Like Lesions (BKL) represents benign skin growths resemble actinic keratoses but have different characteristics. Dermatofibroma (DF) is a benign skin lesion that appears as a firm, round bump and is typically brown or reddish-brown. Melanocytic Nevi (NV) is usually benign and is commonly known as moles.

### 2.3 Ideal Size of Images

The optimal size of images for machine learning algorithms depends on several factors, including the complexity of the model, the computing power at hand, the size and complexity of the dataset, and the application’s specific requirements. Larger images generally contain more detailed information, providing the potential for improved feature extraction and better representation of the image characteristics. ^39,40^ However, employing larger photos requires more memory and computing power to process and analyze the massive amounts of data in the image. Smaller images demand less memory and processing power, making them more suitable for systems with limited resources. However, reducing the image size may lead to the loss of some crucial details or characteristics of the skin lesions, potentially affecting the classification accuracy.^39,41^ To determine the optimal image size, an empirical approach is often adopted. Researchers and developers usually experiment with training and testing the machine learning classification model on images of varying sizes, monitoring performance metrics to determine which image size produces the best results. This empirical exploration allows for a data-driven decision on the ideal image size that balances accuracy, computational efficiency, and memory requirements for the CADx system and dataset.

In this study, we resize the images from 600 x 450 pixels to 64 x 64 pixels. This adjustment was made to expedite training and mitigate the risk of overfitting.

### 2.4 Resampling

Resampling is used to generate a more representative sample of the population, improve the performance of a model, or balance an unbalanced dataset. Resampling is a statistical approach that uses random extract samples from a dataset to build a new dataset with fewer or more samples that have the same distribution as the original dataset. ^42,43^ Resampling is particularly useful when dealing with imbalanced datasets, where one class significantly outnumbers the others, leading to potential biases in the model’s predictions. Table 2 shows the differences between the two types of resampling techniques.

**Table 2:** Two Types of Resampling Techniques.

| Resampling Technique | Key Applications |
| --- | --- |
| Under-sampling | This involves removing some samples from the majority class to balance the minority class |
| Over-sampling | This involves replicating or creating new samples from the minority class to balance the majority class. |

Resampling can assist machine learning models increase their accuracy, especially when the data is imbalanced. However, resampling approaches must be used with caution since they might induce bias and overfitting into the model. To make sure that the resampling procedure does not provide too optimistic findings or deteriorate the model generalizability, proper validation and evaluation of the model performance is required. Resampling offers a practical way to deal with class imbalance and enhance the effectiveness of machine learning models in CADx systems. By utilizing either under-sampling or over-sampling approach, the model can be better equipped to detect patterns and make accurate predictions for both majority and minority classes, ultimately contributing to more robust and reliable skin disease detection and diagnosis.

We apply SMOTE^44–46^ to improve the performance of CADx systems by addressing the class imbalance issues. For the feature vectors of two skin cancer images X1 and X2, SMOTE interpolates between the vectors to generate a new synthetic sample X3. The interpolated sample X3 is derived using the formula shown in Equation 1.

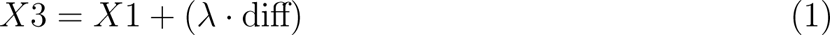

where, 0 *< λ <* 1 and diff = *X*2 *− X*1.

Here, the parameter *λ* controls the extent of interpolation. It is essential to note that the synthetic sample *X*3 is not an actual image from the dataset but rather a new feature vector created through interpolation.

SMOTE increases the representation of minority classes by generating the synthetic samples, thereby offsets the negative consequences associated with skewed datasets. The resampled dataset with synthetic samples can then be used to train the machine learning model, enhancing its ability to generalize and make more accurate predictions for both majority and minority classes. The SMOTE techniques help CADx systems with detecting underrepresented skin lesions.

### 2.5 GAN

GAN operates by concurrently training two neural networks: a generator and a discriminator.^47–50^ The generator is in charge of creating fake images that appear like actual images, while the discriminator is in charge of distinguishing between actual and fake images. During training, the generator learns to make more realistic images, while the discriminator learns to distinguish between actual and fake images, making it more challenging for the generator to produce convincing fakes. GAN models often are used to generate new data that is similar to the original data.

In this work, GAN is used to detect (and eliminate) fake images from the dataset. This is because the existence of fake images can lead to machine learning models learning inaccurate patterns, resulting in poor performance on actual data. By employing GAN, we can distinguish genuine images from potentially misleading ones. Nonetheless, it is critical to recognize that training GANs can be computationally costly and necessitates rigorous hyperparameter tuning. Regardless of this limitation, the use of GAN is expected to improve the dependability and effectiveness of skin cancer classification models. In order to study the significance and effectiveness of GAN, we introduce approximately 25% additional synthetic images for each skin lesion type, amounting to 360 fake images per category in the original HAM10000 dataset.

### 2.6 Machine Learning Techniques

Several machine learning classification models are available to be employed with CADx systems for analyzing images including LDA, SVM, CNN, and an ensemble CNN-SVM.

LDA assumes that the features follow a normal distribution, which can usually be found in medical image data. LDA seeks to enhance the separation between groups, making it particularly effective when there are clear differences between skin lesions. Furthermore, LDA offers a framework for classification, which can be valuable in clinical decision making. However, LDA assumes that classes have the same covariance matrices, which may not always be true in complex medical imaging situations.

SVM is very effective in handling high-dimensional features, which are common in medical image datasets. SVMs can efficiently separate classes even when the data is not linearly separable through the use of different kernel functions, allowing them to capture complex relationships. Additionally, SVMs inherently incorporate the concept of margin, and thus potentially leading to better generalization performance. However, SVMs can be computationally intensive, especially with large datasets, which may impact their real-time applicability in clinical settings.

CNN is capable of handling large, high-resolution images well, making them suitable for skin analysis. CNN excels at extracting the right features from images, eliminating the need for manual feature engineering. This capability is crucial in medical imaging, where intricate patterns and subtle details are vital for accurate diagnosis. Additionally, CNNs are adept at capturing spatial relationships in data, allowing them to discern patterns that may be challenging for traditional machine learning models. However, CNN demands a lot of computing resources, especially when dealing with deep architectures or big data such as the HAM10000 dataset.

Ensemble CNN-SVM models offer a powerful approach for skin cancer classification. This combination leverages the strengths of both models. First, the CNN excels in extracting features from images, capturing complex patterns necessary for accurate diagnosis. Then, the SVM makes explicit separations between classes, which is useful in situations where distinct boundaries are needed. By combining these strengths, the ensemble CNN-SVM model can achieve higher classification accuracy compared to the individual models. Moreover, this approach provides redundancy and robustness against overfitting, as the two models are inherently different in their learning processes and feature extraction methods. However, implementing the ensemble model requires careful tuning and integration, and the computational resources needed for training are substantial. Ensuring compatibility and coherence between the CNN and SVM components is crucial, and finding the right balance between their contributions can be a complex process.

## 3. Machine Learning Models for CADx Systems

In this work, we introduce GAN, EDA, SMOTE, and an ensemble CNN-SVM model in CADx systems to enhance performance of skin neoplasm diagnosis. Figure 3 illustrates the major steps of the proposed CADx system. It should be noted that the preprocessing, segmentation, feature extraction, and feature selection steps are similar to those used in typical CADx systems.

**Figure 3:**
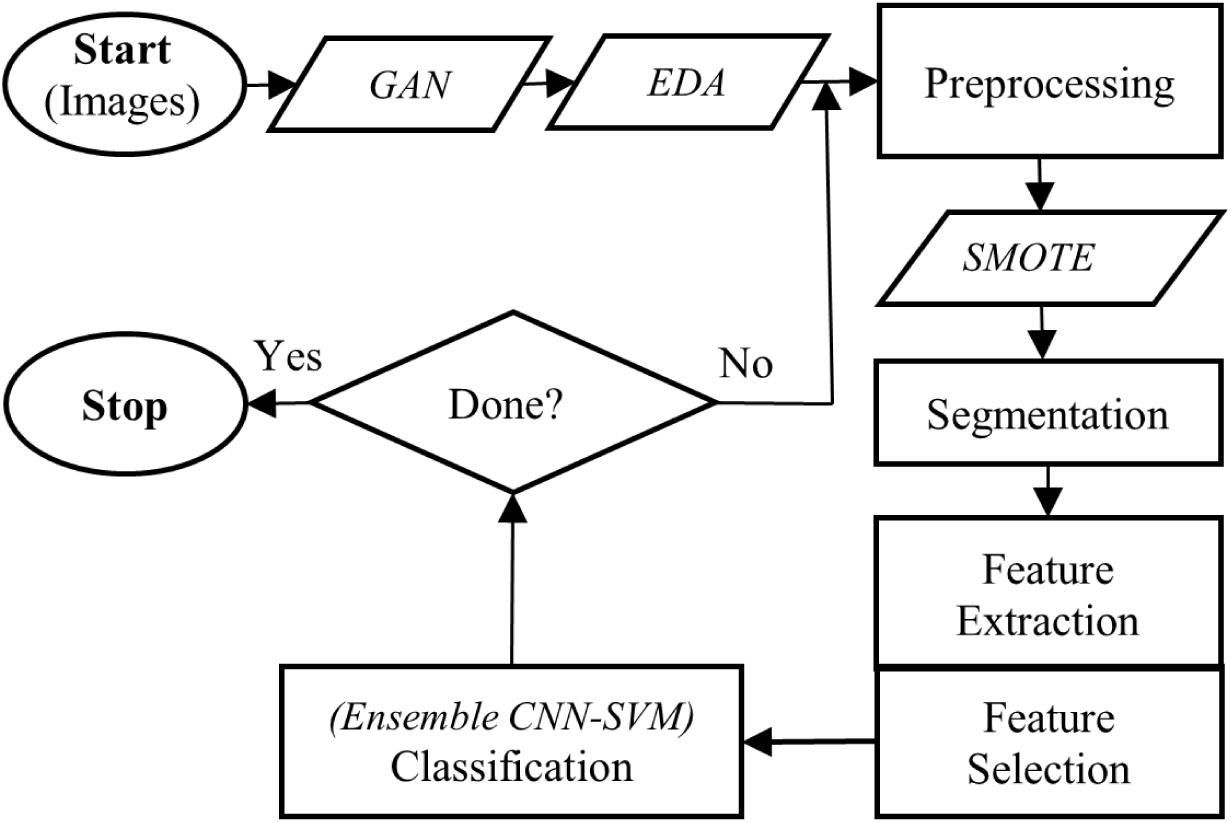
Major Steps of the Proposed CADx System

### 3.1 Application of GAN

In order to tackle concerns regarding the authenticity of the dataset and bolster the reliability of the CADx system, we utilize a GAN as a potent tool for validating the dataset. The GAN is employed to meticulously examine the dataset for any fake images. Through the application of GAN-based analysis, the CADx system can efficiently detect and eliminate any fraudulent samples, ensuring that the model is not trained on deceptive or erroneous data. This validation procedure plays a crucial role in upholding the integrity of the dataset and enhancing the CADx system’s capability to accurately diagnose and classify skin lesions. Consequently, it contributes to the generation of more dependable outcomes in studying dermatological research.

### 3.2 Application of EDA

In order to evaluate and standardize the original dataset to mitigate the risk of machine learning overfitting, we integrate EDA into the proposed CADx system. Beginning with a set of 10,015 images sourced from the HAM10000 dataset, our approach involves augmenting approximately 25% of the dataset with newly generated (fake) images, resulting in a total of 12,535 images. Given that the HAM10000 dataset encompasses seven distinct types, we generate 360 images for each type using a GAN model. The outcomes of the EDA process are depicted in Table 3, illustrating variations among the different types. Incorporating these images into the dataset facilitates the customization and training of machine learning models to effectively discern and accommodate the unique characteristics and patterns exhibited by each category.

**Table 3:** Distribution of the Dataset.

| **Diagnostic Category** | **Original + Fake Images** | **Percentage** |
| --- | --- | --- |
| Melanoma (MEL) | 1473 | 11.75 |
| Basal Cell Carcinoma (BCC) | 874 | 6.97 |
| Vascular Lesion (VAS) | 502 | 4.00 |
| Actinic Keratoses (AKIEC) | 687 | 5.48 |
| Benign Keratosis-Like Lesions (BKL) | 1459 | 11.64 |
| Dermatofibroma (DF) | 475 | 3.79 |
| Melanocytic Nevi (NV) | 7065 | 56.37 |
| Total | 12535 | ~ 100.00 |

### 3.3 Application of SMOTE

Correcting class imbalances within the dataset is imperative for precise skin lesion classification. To address this issue, resampling method SMOTE is utilized. SMOTE generates synthetic samples for the minority class by interpolating existing ones, thereby balancing the dataset. Table 4 showcases the distribution of images within the dataset subsequent to the application of the SMOTE technique. This approach enables the CADx system to learn from a broader and more representative dataset, consequently mitigating overfitting and enhancing generalization capabilities.

**Table 4:**
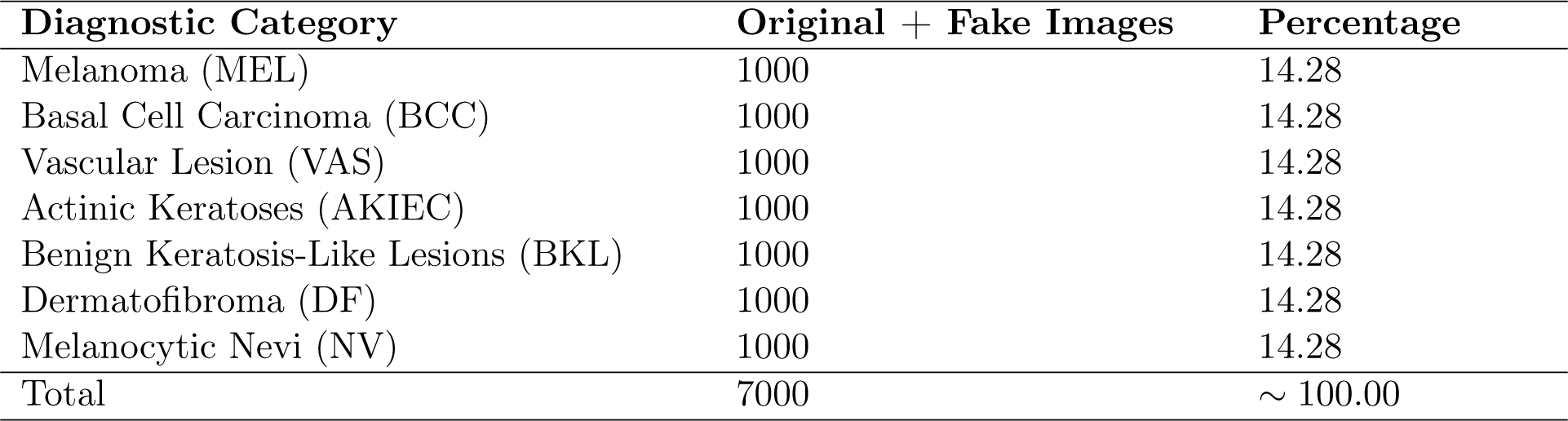
Resampled Dataset using SMOTE.

### 3.4 Ensemble CNN-SVM in Classification

We introduce an ensemble CNN-SVM method in this work to attain enhanced classification accuracy of CADx systems. While SVM excels in managing high-dimensional data, CNN adeptly captures spatial features and hierarchical patterns within images. The ensemble CNN-SVM method harnesses the complementary strengths of CNN and SVM, exploiting both techniques’ prowess to achieve superior generalization and resilience. By implementing these advanced classification techniques, the CADx system enhances its ability to accurately diagnose and classify various skin lesions, providing crucial support to dermatologists and facilitating more effective diagnosis of skin neoplasm.

## 4. Experimental Setup

### 4.1 Dataset Used

In this study, we use 10,015 images from the HAM10000 dataset (in seven types) and assimilate approximately 25% additional synthetic images using GAN. Table 5 shows samples of the seven diagnostic categories of images within the HAM10000 dataset.

**Table 5:**
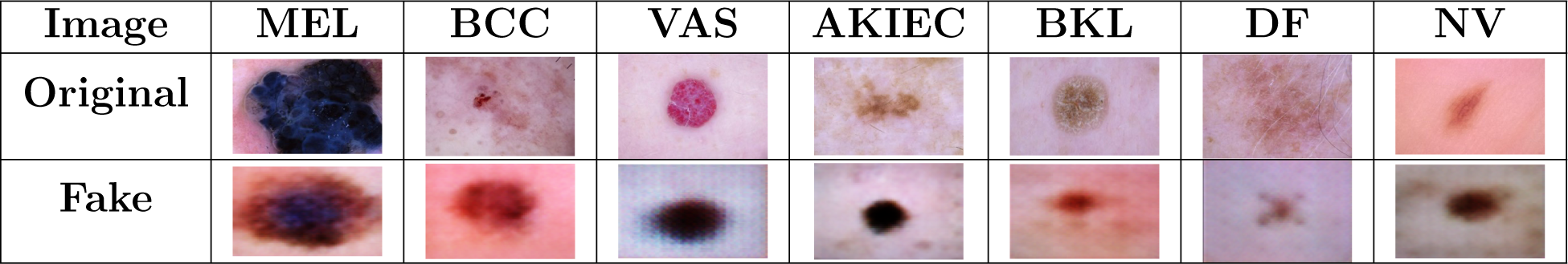
Diagnostic Categories of HAM10000 Dataset.

### 4.2 Model Architecture

In the classification models used in our CADx system, we strategically consider vital hyperparameters to optimize each model’s performance. For LDA, the selection of solver, the application of shrinkage, and the determination of the number of components are pivotal in shaping classification capabilities. Likewise, for SVM, careful parameterization focused on kernel selection, the fine-tuning of the regularization parameter (C), and the degree of the polynomial for optimal decision boundaries are pivotal. The CNN design encompasses key hyperparameters such as the number of convolutional layers, dropout layers, and hidden layers, filter size, max pooling pool size, dropout rate, activation function, L2 (i.e., Euclidean norm) regularization rate, and training epochs. In crafting our ensemble model, we pay significant emphasis on the weighting mechanism for inconsistent predictions to achieve a harmonious fusion of these algorithms. The hyperparameters of the four classification algorithms are summarized in Table 6.

**Table 6:**
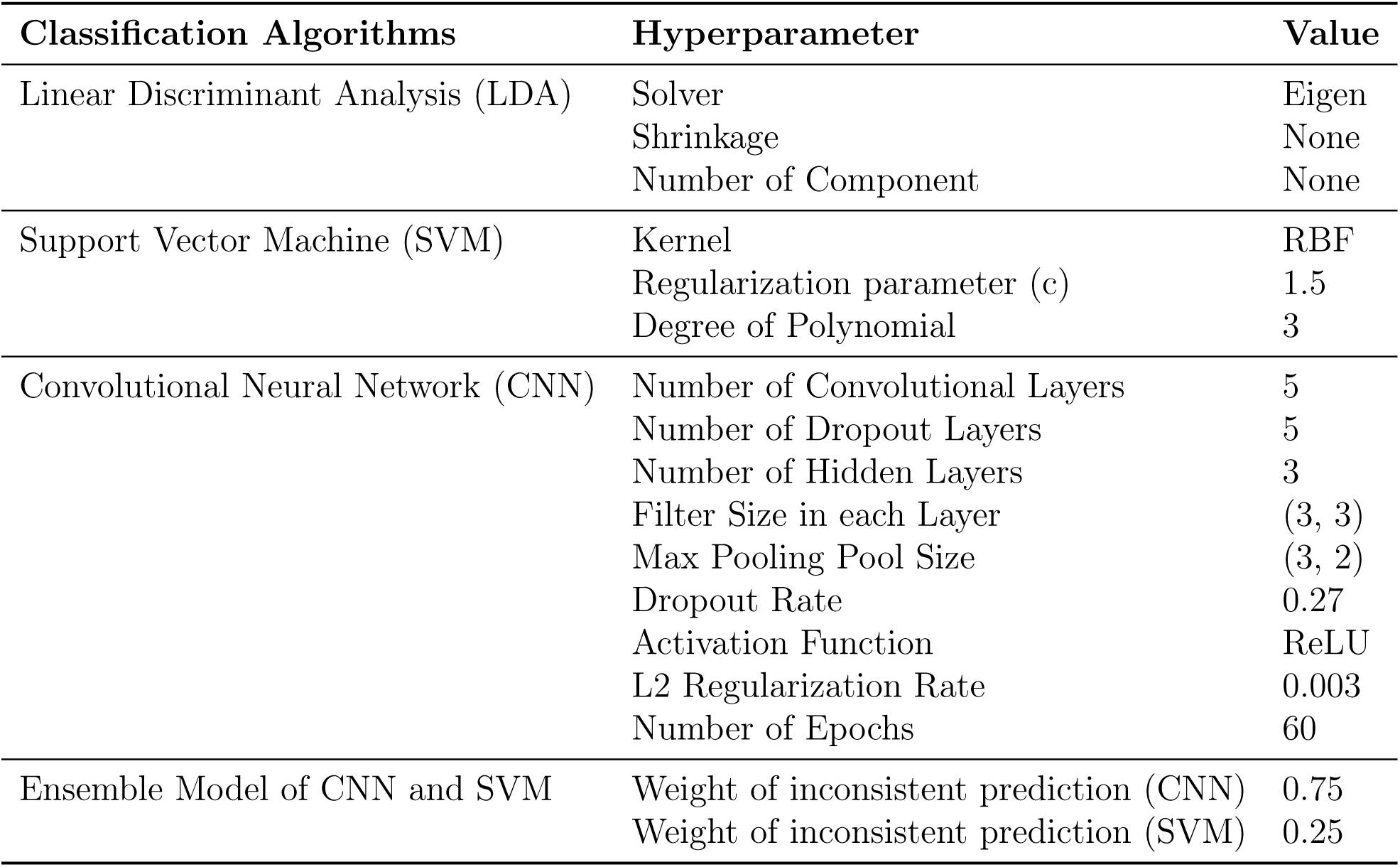
Classification Algorithms and Hyperparameters.

The CNN model consists of five convolutional layers with filter sizes of (3, 3) in each layer, facilitating the extraction of hierarchical features from the input images. After each convolutional layer, max pooling layers with a pool size of (3, 2) are employed to down sample the extracted features, aiding in reducing spatial dimensions. Table 7 shows the shape and number of parameters of the CNN classification algorithm.

**Table 7:** Shape and Number of Parameters of the CNN Model.

| Layer (type) | Output Shape | Param # |
| --- | --- | --- |
| conv2d (Conv2D) | (None, 64, 64, 32) | 896 |
| max_pooling2d (MaxPooling2D) | (None, 32, 32, 32) | 0 |
| dropout (Dropout) | (None, 32, 32, 32) | 0 |
| conv2d_1 (Conv2D) | (None, 32, 32, 64) | 18496 |
| max_pooling2d_1 (MaxPooling2D) | (None, 16, 16, 64) | 0 |
| dropout_1 (Dropout) | (None, 16, 16, 64) | 0 |
| conv2d_2 (Conv2D) | (None, 16, 16, 128) | 73856 |
| max_pooling2d_2 (MaxPooling2D) | (None, 8, 8, 128) | 0 |
| dropout_2 (Dropout) | (None, 8, 8, 128) | 0 |
| conv2d_3 (Conv2D) | (None, 8, 8, 256) | 295168 |
| max_pooling2d_3 (MaxPooling2D) | (None, 4, 4, 256) | 0 |
| dropout_3 (Dropout) | (None, 4, 4, 256) | 0 |
| conv2d_4 (Conv2D) | (None, 4, 4, 512) | 1180160 |
| max_pooling2d_4 (MaxPooling2D) | (None, 2, 2, 512) | 0 |
| dropout_4 (Dropout) | (None, 2, 2, 512) | 0 |
| global_average_pooling2d (GlobalAveragePooling2D) | (None, 256) | 0 |
| dense (Dense) | (None, 128) | 65664 |
| dropout_5 (Dropout) | (None, 128) | 0 |
| dense_1 (Dense) | (None, 7) | 903 |
| Total Parameters |  | 1635143 |

To mitigate overfitting, dropout layers with a dropout rate of 0.27 are strategically inserted after each max pooling layer. The depth of the feature maps increases progressively through the network, starting with 32 filters in the first layer and reaching 256 filters in the final convolutional layer. A global average pooling layer is incorporated to transform the spatial dimensions into a vector of length 256, contributing to reducing the total number of parameters. After the convolutional layers, there are two fully connected dense layers with 127 and 7 units, respectively. The first dense layer employs Rectified Linear Unit (ReLU) activation and L2 regularization with a rate of 0.003, enhancing the model’s ability to learn intricate patterns in data. The entire model comprises a total of 1,635,143 parameters, which include weights and biases. During training, the model undergoes 60 epochs with early stopping and learning rate reduction callbacks, ensuring effective convergence and preventing overfitting.

## 5. Experimental Results

In this section, we discuss experimental results obtained from the proposed CADx system to assess the impact of removing fake data, applying resampled data, and using the ensemble CNN-SVM model. Performance metrics such as precision, recall, F1-score, and accuracy are used for evaluation.

### 5.1 Performance of a Typical CADx System

First, we present performance metrics obtained from a typical CADx system (without using GAN) by utilizing images from the original HAM10000 dataset plus an additional 25% generated/fake images.

The distribution of images among the seven different classes of skin lesions is already summarized in Table 3, shedding light on the representation and prevalence of each skin lesion type. The classification performances of LDA, SVM, and CNN without the implementation of GAN are presented in Tables 8, 9, and 10, respectively.

**Table 8:** LDA Performance without GAN (with Fake Data)

| Category | Precision | Recall | F1-score |
| --- | --- | --- | --- |
| MEL | 0.19 | 0.20 | 0.19 |
| BCC | 0.15 | 0.25 | 0.18 |
| VAS | 0.11 | 0.19 | 0.14 |
| AKIEC | 0.15 | 0.17 | 0.15 |
| BKL | 0.17 | 0.22 | 0.19 |
| DF | 0.00 | 0.00 | 0.00 |
| NV | 0.69 | 0.66 | 0.67 |
| LDA Average | 0.21 | 0.24 | 0.22 |
| LDA Model Accuracy | 0.49 |  |  |

**Table 9:** SVM Performance without GAN (with Fake Data)

| Category | Precision | Recall | F1-score |
| --- | --- | --- | --- |
| MEL | 0.56 | 0.13 | 0.22 |
| BCC | 0.45 | 0.40 | 0.42 |
| VAS | 0.52 | 0.44 | 0.48 |
| AKIEC | 0.36 | 0.14 | 0.20 |
| BKL | 0.55 | 0.29 | 0.38 |
| DF | 0.00 | 0.00 | 0.00 |
| NV | 0.76 | 0.97 | 0.85 |
| SVM Average | 0.45 | 0.33 | 0.36 |
| SVM Model Accuracy | 0.72 |  |  |

The LDA model exhibits notable disparities in its performance across different skin lesion classes (as shown in Table 8). Particularly noteworthy is the poor performance for the DF class, where both precision and recall are reported as 0.00, resulting in an F1-score of0.00. This indicates significant challenges in correctly identifying and classifying instances of DF, highlighting an area where the model may need improvement. In contrast, the model performs relatively well for the NV class, with moderate precision (0.69), recall (0.66) and F1-score (0.67). The model performs poor for the VAS (F1-score 0.14), AKIEC (0.15), BCC (0.18), BKL (0.19), and MEL (0.19) classes, contributing to the model’s average precision of 0.21, average recall of 0.24, average F1-score of 0.22, and a moderate overall accuracy of 0.49.

The SVM model displays varying performance across different skin lesion classes (as shown in Table 9). Notably, it achieves high precision (0.76), recall (0.97), and F1-score (0.85) for the NV class, contributing to the model’s overall accuracy of 0.72. However, the model’s performance varies across other classes, with notable discrepancies in precision, recall, and F1-score, particularly for the DF class, where precision, recall, and F1-score are reported as zero. The average precision (0.45), recall (0.33), and F1-score (0.36) suggest that there are rooms for performance improvement.

The CNN model demonstrates varied performance across different skin lesion classes (as shown in Table 10). Notably, it achieves high precision (0.85), recall (0.94), and F1-score (0.89) for the NV class, contributing to the model’s impressive overall accuracy of 0.77. However, the model’s performance varies across other classes, with some challenges in correctly classifying instances of DF, where the recall is reported as 0.12, resulting in an F1-score of 0.22. The average precision (0.65), recall (0.47), and F1-score 0.52 indicates a reasonable balance between precision and recall across the skin lesion categories.

**Table 10:** CNN Performance without GAN (with Fake Data)

| Category | Precision | Recall | F1-score |
| --- | --- | --- | --- |
| MEL | 0.55 | 0.41 | 0.47 |
| BCC | 0.61 | 0.52 | 0.56 |
| VAS | 0.74 | 0.52 | 0.61 |
| AKIEC | 0.45 | 0.31 | 0.37 |
| BKL | 0.54 | 0.49 | 0.51 |
| DF | 0.80 | 0.12 | 0.22 |
| NV | 0.85 | 0.94 | 0.89 |
| CNN Average | 0.65 | 0.47 | 0.52 |
| CNN Model Accuracy | 0.77 |  |  |

To gain insights into the CNN model’s training process, the key visualization plot of Accuracy versus Epoch (shown in Figure 4) is generated. The observed plateau in accuracy around epoch 35 suggests that the model has reached a saturation point in learning from the training data, indicating that further training beyond this point may not significantly improve performance on the test set. The convergence of training and validation accuracy at 0.86 and 0.77, respectively, indicates that the model generalizes well to unseen data but has reached a point of diminishing returns in learning from the training set.

**Figure 4:**
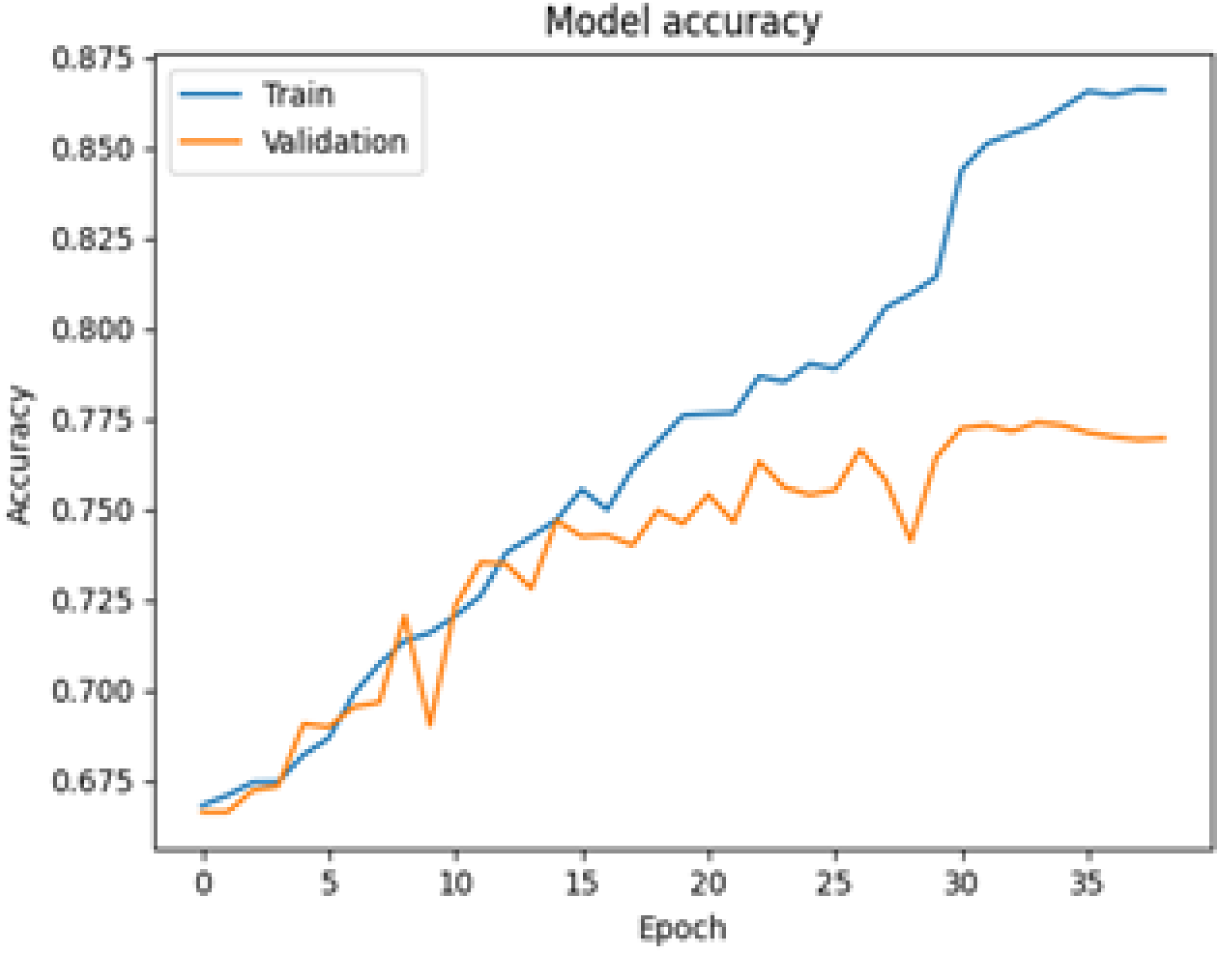
CNN Accuracy versus Epoch without GAN (with Fake Data)

Table 11 shows the performance metrics due to applying the ensemble CNN-SVM model in the CADx system. The ensemble model exhibits notable strengths in accurately classifying the majority of skin lesions, particularly achieving high precision, recall, and F1-score for the NV class with values of 0.97, 0.96, and 0.97, respectively. The results contribute to an impressive overall accuracy of 0.79. However, there are areas for improvement, especially in handling less prevalent classes like AKIEC, where the precision, recall, and F1-score are relatively lower at 0.39, 0.15, and 0.22, respectively.

**Table 11:** Ensemble CNN-SVM Model Performance without GAN (with Fake Data)

| Category | Precision | Recall | F1-score |
| --- | --- | --- | --- |
| MEL | 0.62 | 0.37 | 0.46 |
| BCC | 0.60 | 0.69 | 0.64 |
| VAS | 0.79 | 0.76 | 0.78 |
| AKIEC | 0.39 | 0.15 | 0.22 |
| BKL | 0.56 | 0.62 | 0.59 |
| DF | 0.52 | 0.20 | 0.28 |
| NV | 0.97 | 0.96 | 0.97 |
| CNN-SVM Average | 0.64 | 0.53 | 0.56 |
| CNN-SVM Model Accuracy | 0.79 |  |  |

### 5.2 Performance of the CADx System with GAN

Second, we present performance metrics acquired with a GAN in the CADx system for the original HAM10000 dataset plus 25% generated/fake images. The performance of LDA, SVM, and CNN with GAN is presented in Tables 12, 13, and 14, respectively.

**Table 12:** LDA Performance with GAN (without Fake Data)

| Category | Precision | Recall | F1-score |
| --- | --- | --- | --- |
| MEL | 0.30 | 0.38 | 0.34 |
| BCC | 0.37 | 0.61 | 0.46 |
| VAS | 0.69 | 0.80 | 0.74 |
| AKIEC | 0.44 | 0.64 | 0.52 |
| BKL | 0.38 | 0.39 | 0.38 |
| DF | 0.89 | 0.74 | 0.81 |
| NV | 0.93 | 0.68 | 0.79 |
| LDA Average | 0.57 | 0.60 | 0.58 |
| LDA Model Accuracy | 0.52 |  |  |

**Table 13:** SVM Performance with GAN (without Fake Data)

| Category | Precision | Recall | F1-score |
| --- | --- | --- | --- |
| MEL | 0.56 | 0.48 | 0.51 |
| BCC | 0.69 | 0.80 | 0.74 |
| VAS | 0.89 | 0.93 | 0.90 |
| AKIEC | 0.68 | 0.74 | 0.71 |
| BKL | 0.69 | 0.49 | 0.58 |
| DF | 0.80 | 0.80 | 0.80 |
| NV | 0.85 | 0.97 | 0.91 |
| SVM Average | 0.73 | 0.75 | 0.73 |
| SVM Model Accuracy | 0.76 |  |  |

The LDA model demonstrates relatively balanced performance across various skin lesion classes, as shown in Table 12, achieving notable precision, recall, and F1-score for DF with values of 0.89, 0.74, and 0.81, respectively. However, the model’s overall accuracy is moderate at 0.52. While excelling in certain areas, there are rooms for improvement in enhancing precision, recall, and F1-score for other classes, such as MEL, BKL, and BCC.

The SVM model showcases better performance across various skin lesion classes, as shown in Table 13, achieving particularly high precision, recall, and F1-score for NV with values of 0.85, 0.97, and 0.91, respectively. The model’s overall accuracy is commendable at 0.76, indicating its effectiveness in correctly classifying skin lesions. The balanced precision, recall, and F1-score across multiple classes suggest the model’s capability to generalize well to diverse skin conditions.

The CNN model exhibits promising performance across various skin lesion classes, as shown in Table 14, with notable precision, recall, and F1-score values. Particularly impressive is the perfect precision (1.00), moderate recall (0.76), and high F1-score (0.86) for the DF class, showcasing the model’s ability to accurately identify this specific skin lesion type. The overall accuracy of 0.80 is indicative of the model’s success in correctly classifying skin lesions across diverse categories. The balanced precision, recall, and F1-score across multiple classes emphasize the CNN model’s effectiveness in providing accurate predictions for various skin conditions. The key visualization plot of Accuracy versus Epoch is generated (as shown in Figure 5). The observed stabilization in accuracy is at around epoch 30. With training and validation accuracies converging at 0.87 and 0.80, respectively, the model demonstrates effective generalization to unseen data.

**Figure 5:**
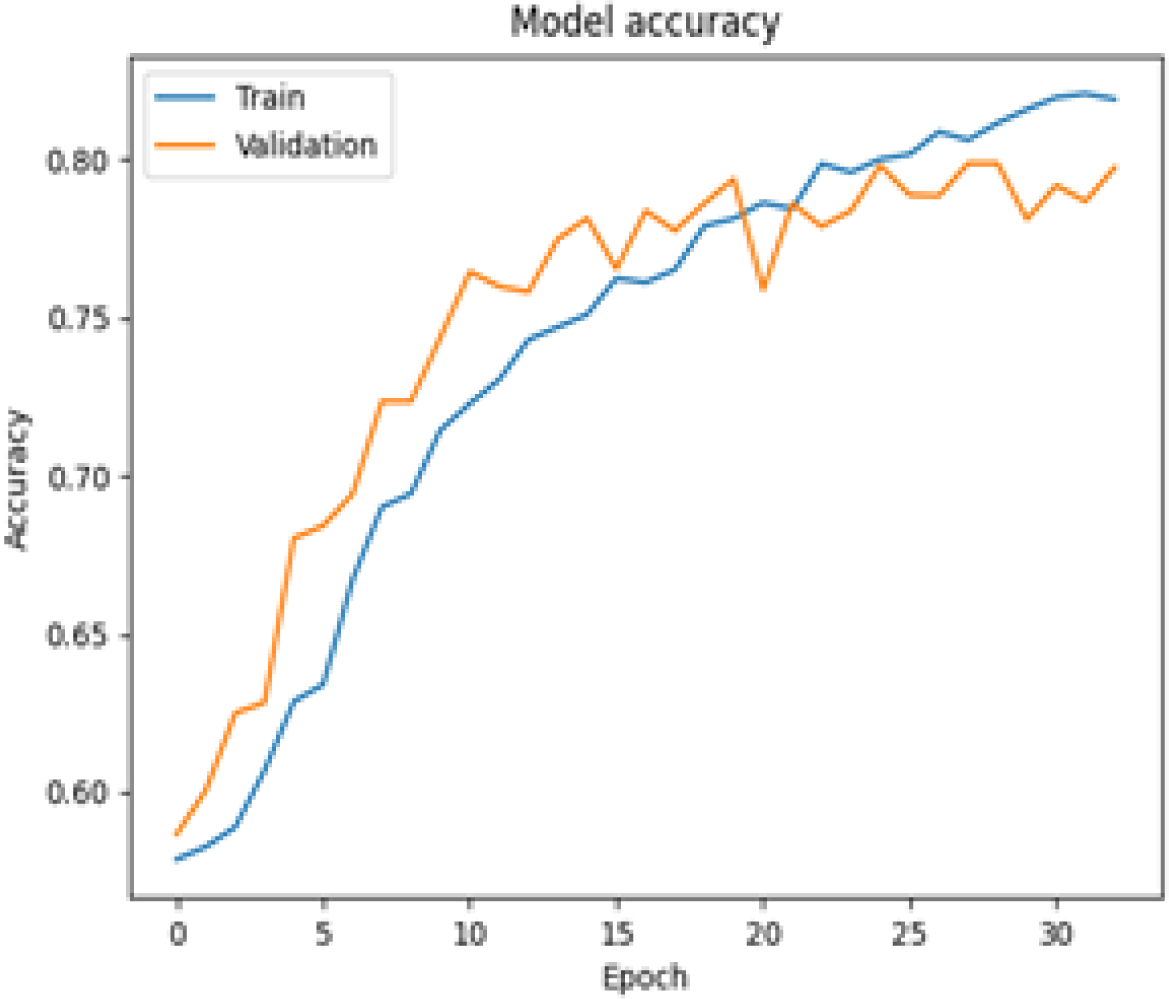
CNN Accuracy versus Epoch with GAN (without Fake Data)

**Table 14:** CNN Performance with GAN (without Fake Data)

| Category | Precision | Recall | F1-score |
| --- | --- | --- | --- |
| MEL | 0.69 | 0.45 | 0.55 |
| BCC | 0.81 | 0.58 | 0.67 |
| VAS | 0.98 | 0.88 | 0.93 |
| AKIEC | 0.87 | 0.64 | 0.74 |
| BKL | 0.74 | 0.51 | 0.60 |
| DF | 1.00 | 0.76 | 0.86 |
| NV | 0.86 | 0.96 | 0.90 |
| CNN Average | 0.85 | 0.68 | 0.75 |
| CNN Model Accuracy | 0.80 |  |  |

The ensemble CNN-SVM model demonstrates strong performance across various skin lesion classes, with precision, recall, and F1-score values indicating its effectiveness in classification. As shown in Table 15, particularly noteworthy is the high precision (0.95), recall (0.95), and F1-score (0.95) for the NV class, showcasing the model’s robust ability to accurately identify this specific skin lesion type. The overall accuracy of 0.82 reflects the model’s success in correctly classifying skin lesions across diverse categories. The balanced precision, recall, and F1-score across multiple classes highlight the ensemble model’s capability to provide accurate predictions for various skin conditions.

**Table 15:** Ensemble CNN-SVM Model Performance with GAN (without Fake Data)

| Category | Precision | Recall | F1-score |
| --- | --- | --- | --- |
| MEL | 0.55 | 0.58 | 0.57 |
| BCC | 0.74 | 0.70 | 0.72 |
| VAS | 0.97 | 0.92 | 0.94 |
| AKIEC | 0.77 | 0.70 | 0.73 |
| BKL | 0.71 | 0.55 | 0.63 |
| DF | 0.97 | 0.87 | 0.92 |
| NV | 0.95 | 0.95 | 0.95 |
| CNN-SVM Average | 0.81 | 0.75 | 0.78 |
| CNN-SVM Model Accuracy | 0.82 |  |  |

### 5.3 Performance Analysis on Resampled Dataset with GAN

Finally, we describe the results collected by resampling the dataset images to 1000 samples for each category, with the use of GAN in the CADx system. The distribution of images among the seven different classes of skin lesions is summarized in Table 4, shedding light on the representation and prevalence of each skin lesion type. The classification performance of LDA, SVM, and CNN for resampled dataset with GAN is presented in Tables 16, 17, and 18, respectively.

**Table 16:** LDA Performance on Resampled Dataset with GAN.

| Category | Precision | Recall | F1-score |
| --- | --- | --- | --- |
| MEL | 0.64 | 0.66 | 0.65 |
| BCC | 0.73 | 0.70 | 0.71 |
| VAS | 0.93 | 0.98 | 0.85 |
| AKIEC | 0.76 | 0.80 | 0.78 |
| BKL | 0.61 | 0.59 | 0.60 |
| DF | 0.91 | 0.92 | 0.91 |
| NV | 0.75 | 0.68 | 0.71 |
| LDA Average | 0.76 | 0.76 | 0.76 |
| LDA Model Accuracy | 0.76 |  |  |

**Table 17:** SVM Performance on Resampled Dataset with GAN.

| Category | Precision | Recall | F1-score |
| --- | --- | --- | --- |
| MEL | 0.69 | 0.69 | 0.70 |
| BCC | 0.79 | 0.89 | 0.84 |
| VAS | 0.97 | 1.00 | 0.98 |
| AKIEC | 0.91 | 0.91 | 0.92 |
| BKL | 0.77 | 0.67 | 0.72 |
| DF | 0.94 | 0.98 | 0.96 |
| NV | 0.81 | 0.71 | 0.83 |
| SVM Average | 0.84 | 0.85 | 0.85 |
| SVM Model Accuracy | 0.83 |  |  |

The LDA model exhibits notable performance in classifying skin lesions, with high precision, recall, and F1-score values across various classes as shown in Table 16. For LDA model, particularly remarkable is the model’s proficiency in identifying the DF class, achieving precision, recall, and F1-score of 0.91, 0.92, and 0.91, respectively. The overall accuracy of 0.76 underscores the model’s success in providing accurate predictions for different skin lesion types. The balanced precision, recall, and F1-score across multiple classes highlight the LDA model’s capability to handle diverse skin conditions effectively.

The SVM model demonstrates exceptional performance in skin lesion classification (as shown in Table 17), with high precision, recall, and F1-score values across various classes. Particularly noteworthy is the model’s accuracy in distinguishing the VAS class, achieving precision, recall, and F1-score of 0.97, 1.00, and 0.98, respectively. The overall accuracy of 0.83 underscores the model’s success in providing precise and reliable predictions for different skin lesion types. The balanced precision and recall scores across multiple classes highlight the SVM model’s effectiveness in handling diverse skin conditions.

Table 18 exhibits outstanding performance of the CNN model in skin lesion classification, demonstrating high precision, recall, and F1-score values across different classes. Notably, the model achieves exceptional accuracy in recognizing the DF class, with precision, recall, and F1-score reaching 0.99, 1.00, and 0.99, respectively. The overall accuracy of 0.87 reflects the model’s excellence in providing accurate predictions for various skin lesion types. Balanced precision, recall, and F1-score across multiple classes underscore the CNN model’s robustness and efficacy in handling diverse skin conditions.

**Table 18:** CNN Performance on Resampled Dataset with GAN.

| Category | Precision | Recall | F1-score |
| --- | --- | --- | --- |
| MEL | 0.77 | 0.74 | 0.75 |
| BCC | 0.90 | 0.90 | 0.90 |
| VAS | 0.96 | 1.00 | 0.98 |
| AKIEC | 0.92 | 0.92 | 0.92 |
| BKL | 0.77 | 0.74 | 0.75 |
| DF | 0.99 | 1.00 | 0.99 |
| NV | 0.78 | 0.79 | 0.78 |
| CNN Average | 0.87 | 0.87 | 0.87 |
| CNN Model Accuracy | 0.87 |  |  |

The key visualization plot of Accuracy versus Epoch is shown in Figure 6. The observed stabilization in accuracy is at around epoch 32. With training and validation accuracy converging at 0.91 and 0.83, respectively, the model demonstrates effective generalization to unseen data.

**Figure 6:**
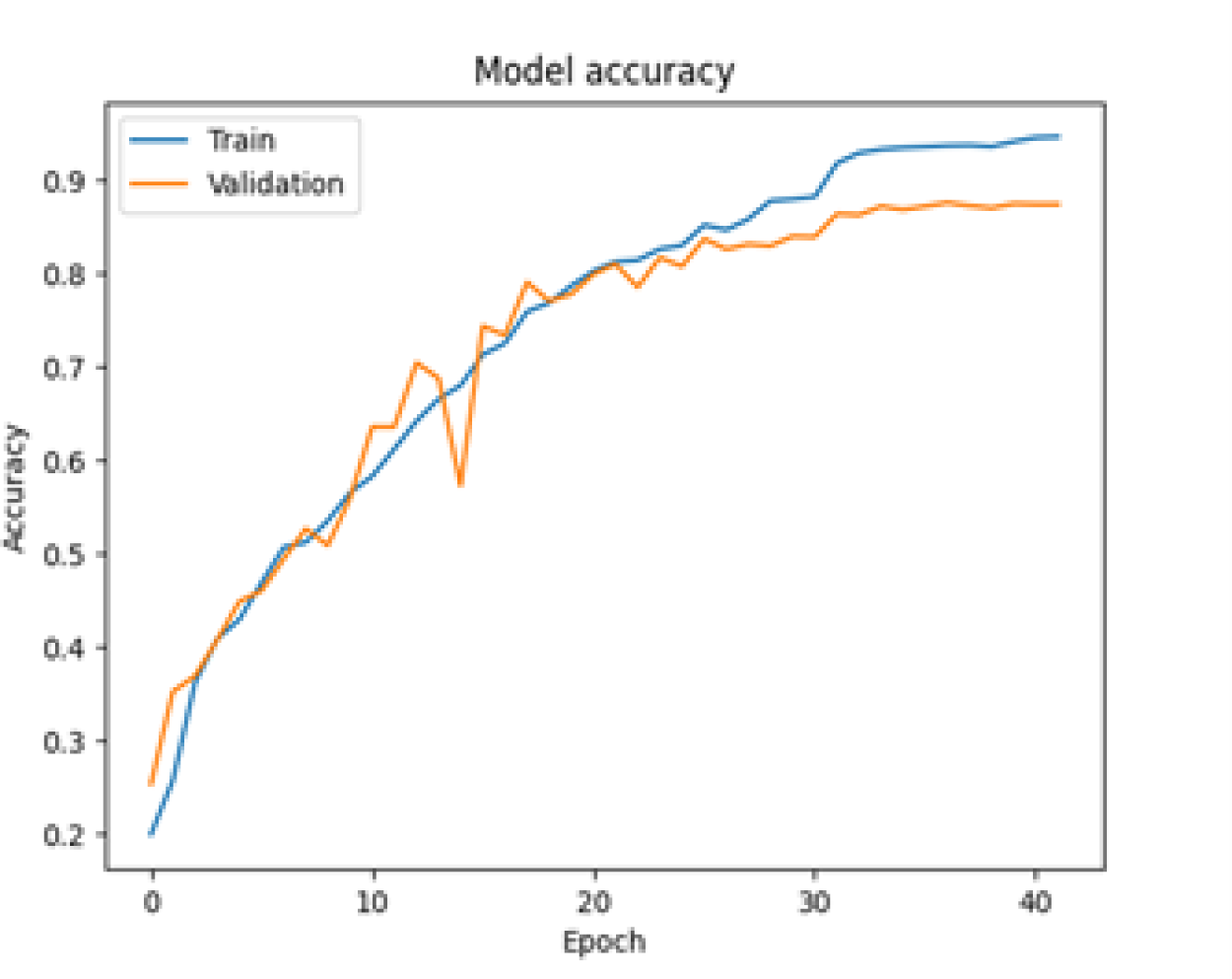
CNN Accuracy vs Epoch for Resampled Dataset with GAN

The ensemble CNN-SVM model, combining the strengths of CNN and SVM, showcases outstanding performance in skin lesion classification (as shown in Table 19). With precision, recall, and F1-score values consistently above 0.90 for most classes, the model demonstrates a high level of accuracy in identifying various skin lesion types. Particularly noteworthy is the exceptional performance in recognizing the VAS class, with precision, recall, and F1-score reaching 0.99, 1.00, and 0.99, respectively. The overall accuracy of 0.94 highlights the effectiveness of the ensemble model in providing accurate predictions for diverse skin conditions, making it a robust and reliable tool for skin lesion diagnosis. The balanced precision, recall, and F1-score across multiple classes further emphasize the model’s ability to handle different skin conditions with high accuracy.

**Table 19:** Ensemble CNN-SVM Model Performance on Resampled Dataset with GAN.

| Category | Precision | Recall | F1-score |
| --- | --- | --- | --- |
| MEL | 0.79 | 0.79 | 0.79 |
| BCC | 0.92 | 0.96 | 0.94 |
| VAS | 0.99 | 1.00 | 0.99 |
| AKIEC | 0.95 | 0.97 | 0.96 |
| BKL | 0.90 | 0.78 | 0.84 |
| DF | 0.97 | 1.00 | 0.98 |
| NV | 0.89 | 0.97 | 0.93 |
| CNN-SVM Average | 0.92 | 0.93 | 0.92 |
| CNN-SVM Model Accuracy | 0.94 |  |  |

### 5.4 Summary of Experimental Results

Accuracy and F1-scores of LDA, SVM, CNN, and ensemble model of CNN and SVM is summarized in Table 20. The utilization of a resampled dataset, with the incorporation of GAN, has demonstrated substantial improvement in skin lesion classification across various models. The enhancement in accuracy is notably significant, emphasizing the effectiveness of addressing class imbalance through resampling techniques. LDA initially exhibits the lowest accuracy, followed by SVM and CNN, while the ensemble model of CNN and SVM consistently delivers the highest accuracy.

**Table 20:** Summary of Accuracy and F1-score.

| **Classification Model** | **Accuracy and F1 Score** | **Dataset without GAN** | **Dataset with GAN** | **Resampled Dataset with GAN** |
| --- | --- | --- | --- | --- |
| LDA | Accuracy | 0.49 | 0.52 | 0.76 |
|  | F1 Score | 0.22 | 0.58 | 0.76 |
| SVM | Accuracy | 0.72 | 0.76 | 0.83 |
|  | F1 Score | 0.36 | 0.73 | 0.85 |
| CNN | Accuracy | 0.77 | 0.80 | 0.87 |
|  | F1 Score | 0.52 | 0.75 | 0.87 |
| CNN-SVM | Accuracy | 0.79 | 0.78 | 0.94 |
|  | F1 Score | 0.48 | 0.82 | 0.92 |

The introduction of GAN effectively mitigates issues associated with fake images, enhancing overall accuracy, precision, recall, and F1-score across all classification algorithms. Furthermore, the resampled dataset (with GAN) consistently outperforms the dataset with and without GAN, achieving the highest accuracy across all models. This highlights the effectiveness of resampling in addressing class imbalance, allowing the models to better generalize and make accurate predictions for all skin lesion classes. The combination of GAN, EDA, and resampling techniques significantly improves the performance of the classification models, emphasizing the importance of dataset quality and balance in achieving accurate and reliable results.

## Conclusions

Skin lesion classification is a pivotal aspect of dermatological diagnostics, and the development of effective CADx systems is imperative for accurate and timely detection of skin diseases. This study addresses the challenges inherent in skin lesion classification by introducing novel techniques such as GAN for fake image detection, resampling methods, and the implementation of an ensemble model combining CNN and SVM. Our contributions aim to enhance the robustness and accuracy of CADx systems, laying the groundwork for more reliable dermatological diagnostics.

The experimental results show significant advancements across various classification models, particularly when employing a GAN with resampled dataset. The notable increase in accuracy underscores the efficacy of mitigating class imbalance through resampling techniques. LDA initially demonstrates the lowest accuracy (0.49), followed by SVM (0.72), CNN (0.77), and ensemble CNN-SVM model (0.79). The incorporation of GAN effectively addresses challenges associated with fake images, leading to improvements in overall accuracy, precision, recall, and F1-score across all classification algorithms. The resampled dataset, augmented with GAN, consistently outperformed the original dataset, emphasizing the critical role of addressing class imbalance in facilitating accurate predictions for all skin lesion classes. Finally, LDA exhibits a moderate accuracy of (0.76), followed by SVM (0.83), CNN (0.87), and ensemble CNN-SVM model (0.94, the highest accuracy).

We plan to explore the integration of advanced machine learning techniques, the exploration of additional data processing methods, and the development of more sophisticated ensemble models in our future endeavor.

